# Real-Time Sensing of Food Spoilage Using a Recombinant mVenus–Tolles FRET pH Biosensor

**DOI:** 10.64898/2025.11.30.691354

**Authors:** Matan Gabay, Marina Sova, Tal Laviv, Maayan Gal

## Abstract

Monitoring food spoilage is essential for enhancing food safety and reducing waste. pH changes serve as a valuable indicator of microbial activity, and real-time pH monitoring can provide an accurate and non-invasive indication of food spoilage. The pHlameleon chimera proteins were developed for pH sensing by Förster resonance energy transfer (FRET) and were extensively used in various biomedical applications. Herein, we evaluate the mVenus–Tolles as a FRET-based biosensor for detecting pH changes as a proxy for food spoilage. The protein was fused to an N-terminal vesicle-nucleating peptide (VNP) tag and recombinantly expressed and purified to homogeneity. Experimental validation demonstrated pH-responsive FRET signal in an array of buffers as well as in a complex food matrix such as chickpea paste, correlating with increasing acidity and microbial growth in food. These findings suggest that this protein-based FRET biosensor holds promise for safe integration into food or packaging for real-time freshness monitoring.

## 1. Introduction

Food spoilage poses major health risks and is considered a significant economic burden (Bhatia, Jha, Sarkar, & Sarangi, 2023; Campoy-Munoz, Cardenete, Delgado, & Sancho, 2021; de Gorter, Drabik, Just, Reynolds, & Sethi, 2021; Snyder, Martin, & Wiedmann, 2024). During food storage, microbial proliferation and enzymatic activity can lead to the formation of off-flavors and harmful toxins, often resulting in foodborne illness (Duchenne, Ranghoo-Sanmukhiya, & Neetoo, 2021; Faour-Klingbeil & E, 2019). Moreover, undetected spoilage not only compromises safety but also accelerates food waste and therefore, methods for monitoring spoilage processes are essential for ensuring food safety and extending shelf life.

Microorganism growth and metabolism within the food matrices could lead to an alkaline environment due to protein degradation or an acidic environment owing to fermentation-based organic acids production (Galvez, Abriouel, Benomar, & Lucas, 2010;J. Zhang, Liu, Xie, Wang, Zhong, & Fan, 2024). As such, pH serves as a reliable and sensitive indicator of microbial growth and spoilage in food matrices. Given its importance, numerous strategies have been developed to monitor pH in food, offering alternatives to traditional electrode-based pH meters (Steinegger, Wolfbeis, & Borisov, 2020). These approaches include natural or synthetic bio-packaging with embedded pH-indicator films (Luo et al., 2023; J. Zhang, Liu, et al., 2024), wireless electronic sensing systems (Shrestha, Kim, Jung, Kim, Truong, & Cho, 2021) and radio-frequency technologies for monitoring the pH (Huang, Deb, Seo, Rao, Chiao, & Chiao, 2012). Additional approaches include pH-responsive injectable bio-ink (Xu et al., 2021), pH-sensitive hydrogels (Diana et al., 2024) and, most commonly, colorimetric dyes operating in the visible or infrared spectrum (Chalitangkoon & Monvisade, 2021; Elhadef et al., 2024; He et al., 2023; J. Liu et al., 2019; Tavassoli, Alizadeh Sani, Khezerlou, Ehsani, Jahed-Khaniki, & McClements, 2022; J. Wang et al., 2024).

In addition to the colorimetric methods, a variety of fluorescence probes have been developed. Fluorescence readout exhibits superior quantitative precision and resolution, enabling better detection of subtle pH fluctuations in food. They are also less susceptible to interference from sample turbidity or the presence of native food chromophores. Indeed, a large array of fluorescence probes for sensing pH has been successfully developed and applied for pH sensing in food systems (Ezati, Khan, Rhim, Kim, & Molaei, 2023; Jiang, Ye, Ma, Rodrigues, Sheng, & Min, 2022; Y. Liu, Yu, et al., 2022; Y. Liu, Zhang, et al., 2022; Sandeep et al., 2024;L. Wang, Xin, Zhang, Ran, Tang, & Cao, 2021; Xiao, Shen, Cao, & Sun, 2023; Yan et al., 2023; Zeng, Xiao, Ye, Ma, & Zhou, 2022;J. Zhang, Yang, Zeng, & Wang, 2024;Y. Zhang et al., 2021; Zhu, Jin, Gao, Gui, & Wang, 2017). A subgroup of fluorescent-based methods is Förster resonance energy transfer (FRET), occurring when an excited fluorescent donor transfers energy to a nearby acceptor fluorophore, with the efficiency of this transfer being highly dependent on the spatial distance between the donor-acceptor pair (Piston & Kremers, 2007; Sekar & Periasamy, 2003;X. Zhang et al., 2019). The distinct spectral signature of FRET reduces background autofluorescence and interference with other fluorophores, allowing for enhanced sensitivity and more accurate signal quantification. These advantages have advanced multiple FRET-based approaches for pH sensing in food (Borchert, Kerry, & Papkovsky, 2013; Li et al., 2025; Xie et al., 2024; G. Zhang et al., 2019; Zhao, Xiao, Tang, Zhou, Qin, & Wang, 2025; Zlotnikov, Savchenko, & Kudryashova, 2023).

Most of the aforementioned fluorescence and FRET pH sensors rely on synthetic or naturally-derived small molecules. In this study, we explored an alternative approach using a recombinant protein-based FRET biosensor to monitor pH changes in food. Specifically, we focused on a member of the pHlameleon family, which are FRET-based protein sensors originally designed for intracellular pH imaging (Burgstaller et al., 2019; Esposito, Gralle, Dani, Lange, & Wouters, 2008). These constructs typically fuse a pH-insensitive cyan fluorescent protein (CFP) donor with a pH-sensitive yellow fluorescent protein (YFP) acceptor. One such design, created to monitor autophagy, involves the fusion of the CFP variant Tolles to YPet, producing a robust pH-responsive biosensor. Such a FRET pair was developed to monitor cellular autophagy by fusing the CFP protein Tolles to YPet, forming a robust pH-sensitive biosensor (Katayama et al., 2020; Nguyen & Daugherty, 2005). Herein, we engineered a related construct by fusing mVenus to Tolles, aiming to evaluate its suitability as a biosensing probe for tracking pH in a complex food environment.

## 2. Materials and Methods

### 2.1 Cloning of the mVenus-Tolles construct

The gene encoding the Tolles fluorescent protein was synthesized by Invitrogen (Thermo Fisher Scientific) based on the sequence reported by Katayama et al. (Katayama et al., 2020). The mVenus gene was obtained as a gift from Steven Vogel (Addgene plasmid #27794) (Koushik, Chen, Thaler, Puhl, & Vogel, 2006). Both the Tolles and mVenus open reading frames (ORFs) were cloned into a plasmid containing the vesicle nucleating peptide (VNP) sequence, gifted by Dan Mulvihill (Addgene plasmid #182386) (Eastwood et al., 2023) using Gibson Assembly. The ORF of Tolles was PCR-amplified with 5’ and 3’ overhangs designed for in-frame fusion to the flexible linker. Similarly, the mVenus ORF was amplified with overhangs complementary to the linker and downstream vector regions. Gibson Assembly was performed according to the manufacturer’s instructions (NEBuilder HiFi DNA Assembly Master Mix, New England Biolabs). The final construct encoded a fusion protein of mVenus– linker–Tolles with N-terminus vnp secretion signal. Primers and construct sequence are listed in supplementary information.

### 2.2 Structural modeling and visualization

The three-dimensional structure of the mVenus–Tolles FRET biosensor was predicted using AlphaFold3 (Abramson et al., 2024).Structural visualization and image generation were performed using UCSF ChimeraX (version 1.9) (Pettersen et al., 2021).

### 2.3 Protein expression and purification

The gene encoding the full-length mVenues-Tolles was cloned into the VNP-containing plasmid as previously described (Eastwood et al., 2023). The resulting plasmid was transformed into *E. coli* BL21 (DE3) cells for recombinant protein expression. Cultures were grown in LB medium at 37 °C with shaking (200 rpm) until the optical density at 600 nm reached approximately 0.8. Protein expression was induced with 1 mM isopropyl β-D-1-thiogalactopyranoside (IPTG), followed by incubation at 22 °C for 16 hours. Cells were harvested and lysed by four cycles of 2-minute sonication, with cooling intervals, then centrifuged at 17,000 × g for 30 minutes. The resulting supernatant was applied to a Ni^2+^-NTA affinity chromatography column, washed with buffer containing 50 mM Tris-HCl (pH 8.0), 300 mM NaCl, and 60 mM imidazole. The Hisx6-tagged protein was eluted using 300 mM imidazole. Eluted fractions were analyzed by 12% SDS–PAGE to verify purity.

### 2.4 FRET measurements and analysis

FRET measurements were conducted using a multimode plate reader (Synergy H1, BioTek). The donor fluorophore (Tolles) was excited at 405 nm, and its emission was recorded at 500 nm. The acceptor fluorophore (mVenus) was excited at 515 nm, with emission detected at 530 nm. FRET was measured using donor excitation (405 nm) and acceptor emission (530 nm). The corrected FRET intensity was calculated using the equation: FRET(corrected) = I(DA) − α·I(DD) − β·I(AA), where: I(DA) is the measured fluorescence at donor excitation and acceptor emission (405/530 nm), I(DD) is donor-only emission at 405/500 nm, I(AA) is acceptor-only emission at 515/530 nm, α accounts for donor spectral bleed-through and is calculated as the ratio I(DA)/I(DD) for donor-only samples, β accounts for acceptor cross-excitation and is calculated as I(DA)/I(AA) for acceptor-only samples.

### 2.5 FRET Assay Across a pH Gradient

To study the pH sensitivity of the mVenus–Tolles FRET biosensor, fluorescence measurements were performed across a broad pH range using purified protein. Buffer solutions covering pH values of 2, 3, 4, 5, 7, 10, 11, and 13 were prepared using standard buffers, and pH was measured using an electrode-based pH meter. For each pH condition, 200 μL of the respective buffer was dispensed into wells of a black 96-well microplate. Donor-only, acceptor-only, and full biosensor constructs were added to the wells in triplicate in a final concentration of 5 μM per well. FRET signal was recorded as described previously, enabling characterization of the biosensor’s pH-dependent response profile.

### 2.6 Preparation of food samples

Hummus samples were prepared by blending 60 g of pre-cooked chickpeas with 100 mL of double-distilled water (DDW) at room temperature using a standard laboratory blender. The resulting homogenized paste was used immediately in downstream experiments to ensure consistency and freshness.

### 2.7 Bacterial counting

To quantify microbial growth, hummus samples were serially diluted in sterile Luria broth (LB) and plated on MRS agar (recipe in supplementary data) using the spread plate technique. Plates were incubated at 37 °C for 16–18 hours, after which visible colonies were counted manually for each dilution. Bacterial load was finalized as colony-forming units per milliliter (CFU/mL). Each measurement was performed in triplicate to ensure reproducibility.

### 2.8 pH measurements

The pH of hummus samples was assessed using colorimetric pH indicator strips: MQuant 5– 10 (Merck, catalog #109533) for moderate pH shifts and MQuant 0–14 (Merck, catalog #109535) for broader range detection. Strips were briefly immersed in the sample surface, and pH values were determined by visual comparison against the manufacturer’s color chart.

## 3. Results

### 3.1 Design and structural modeling of the protein FRET biosensor

To enable direct pH sensing in food, we expressed and purified a chimeric protein composed of the yellow fluorescent protein mVenus and the cyan fluorescent protein Tolles (mVenus– Tolles). Initial expression in *E. coli*, even in the presence of a robust solubility tag such as the maltose binding protein (MBP), yielded predominantly insoluble protein **(Figure S1)**. To improve solubility, we engineered a modified construct incorporating an N-terminal vesicle nucleating peptide (VNP) tag, previously shown to enhance soluble protein expression in bacteria **(Figure 1A and Figure S1)** (Eastwood et al., 2023). A C-terminal Hisx6 tag was added to facilitate affinity purification. The two fluorescent domains were connected via a flexible glycine- and serine-rich linker (ELGGGGSGGGGSGGGGSLE), permitting proper folding and spatial orientation of the two protein fluorophores. Structure generation by AlphaFold and modeling of the chimera protein confirmed that the donor and acceptor domains were correctly oriented and unaffected by the VNP fusion. Moreover, the construct forms a compact configuration, with mVenus (yellow) and Tolles (purple) positioned within 5 nm of each other, well within the FRET-effective range (**Figure 1B**). This architecture supports the potential of the recombinant construct to function as an efficient FRET biosensor for monitoring pH.

**Figure 1.**
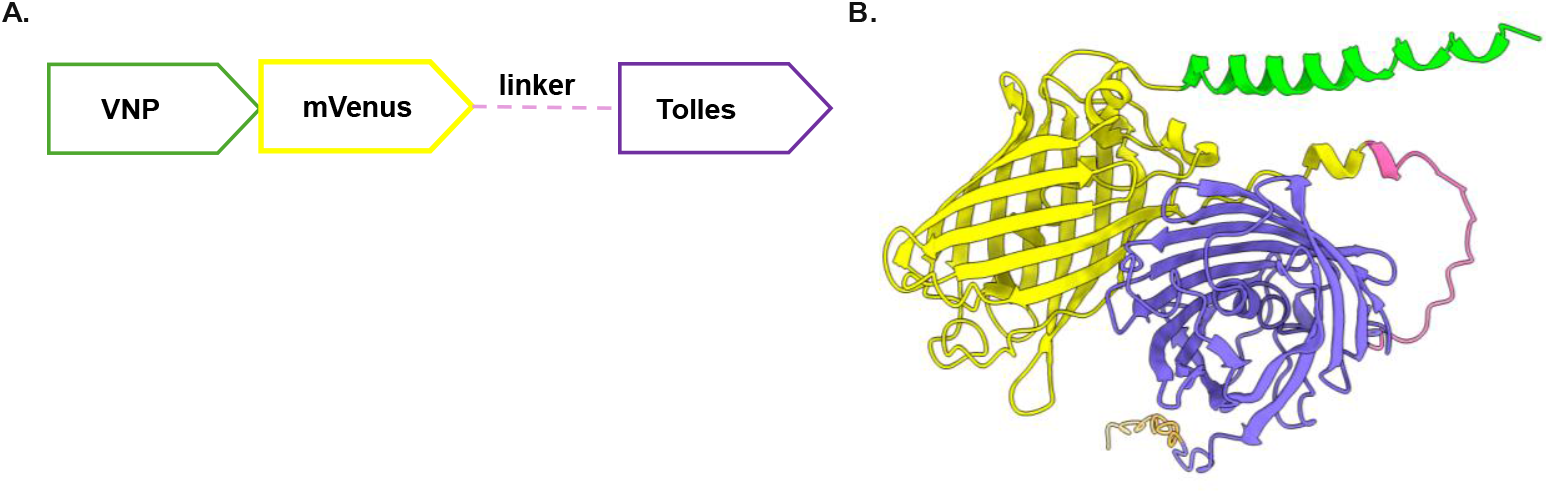
Design and structural modeling of the mVenus–Tolles recombinant FRET biosensor. **(A)** Illustration of the recombinant protein construct containing an N-terminal VNP solubility tag, and the two fluorescent proteins: Tolles (donor) and mVenus (acceptor), separated by a flexible glycine/serine-rich linker. (B) Predicted 3D structure showing the spatial arrangement of domains: VNP (green), mVenus (yellow), linker (pink), Tolles (purple), and Hisx6-tag (orange). The proximity of the fluorophores supports efficient FRET energy transfer.

### 3.2 mVenus-Tolles protein expression and purification

The incorporation of the N-terminal VNP tag facilitated the development of an efficient protocol for recombinant expression and purification of the mVenus–Tolles biosensor. The expression plasmid encoding the protein was transformed into *E. coli* BL21 (DE3) cells. Bacteria were cultured at 37 °C until reaching an optical density (OD_600_) of 0.8, after which protein expression was induced by adding 0.5 mM IPTG, followed by incubation at 25 °C for 16 hours. Cells were subsequently harvested, and the soluble fraction containing the recombinant fusion protein was loaded onto a nickel-affinity chromatography column. Non-specifically bound contaminants were removed using wash buffer containing 40 mM imidazole, and the recombinant biosensor protein was eluted using 250 mM imidazole, resulting in a highly purified chimera protein (Figure 2).

**Figure 2.**
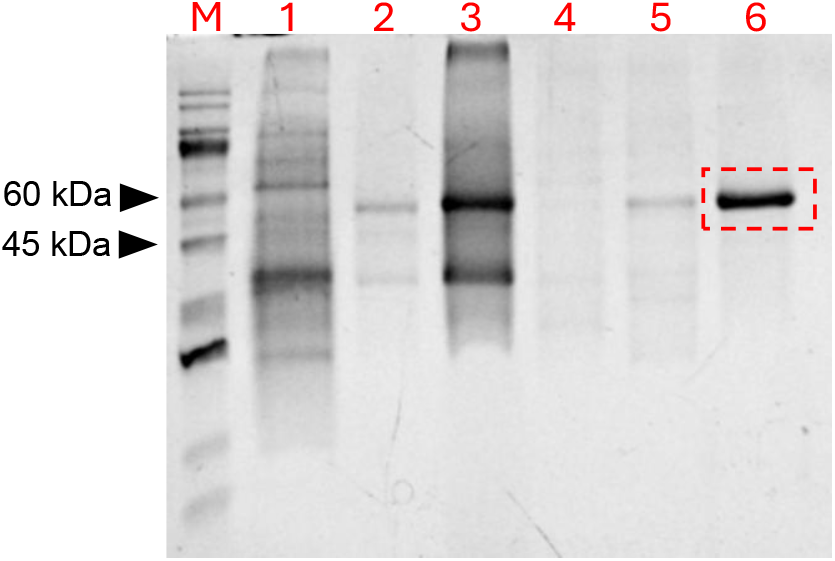
Purification of the mVenus–Tolles recombinant chimera protein. SDS-PAGE gel showing protein samples corresponding to the purification steps: 1 – bacterial cell lysate before induction; 2 – insoluble cell pellet fraction after lysis; 3 – soluble fraction (clarified cell lysate, column load); 4 – column flow-through (unbound fraction); 5 – column wash fraction with 40 mM imidazole; 6 – purified biosensor protein eluted with 250 mM imidazole, with a molecular weight of 58.6 kDa.

### 3.3 FRET response to pH in buffer and food

We next evaluated the FRET response of the recombinant mVenus–Tolles pH sensor in both buffer and food matrix, both under predefined pH conditions. As an initial step, we measured the FRET signal of the sensor at varying concentrations in phosphate-buffered saline (PBS), revealing more than a three-order-of-magnitude signal-to-noise ratio at a concentration above 3 μM (Figure 3A). The protein was then incubated in buffers spanning a range of pH values at a fixed concentration of 5 μM, t. The resulting normalized corrected FRET signal displayed clear pH sensitivity, particularly across the range of pH 4–7, which is the most relevant range for food spoilage detection (Figure 3B). To assess the biosensor performance in a real food matrix, the biosensor was added to chickpea paste (hummus) samples that were adjusted to defined pH values. The FRET pattern in these samples closely mirrored the buffer-based results, confirming that the sensor retains pH responsiveness in a complex food environment (Figure 3C). These results support the feasibility of using mVenus–Tolles as a functional pH biosensor in real food products.

**Figure 3.**
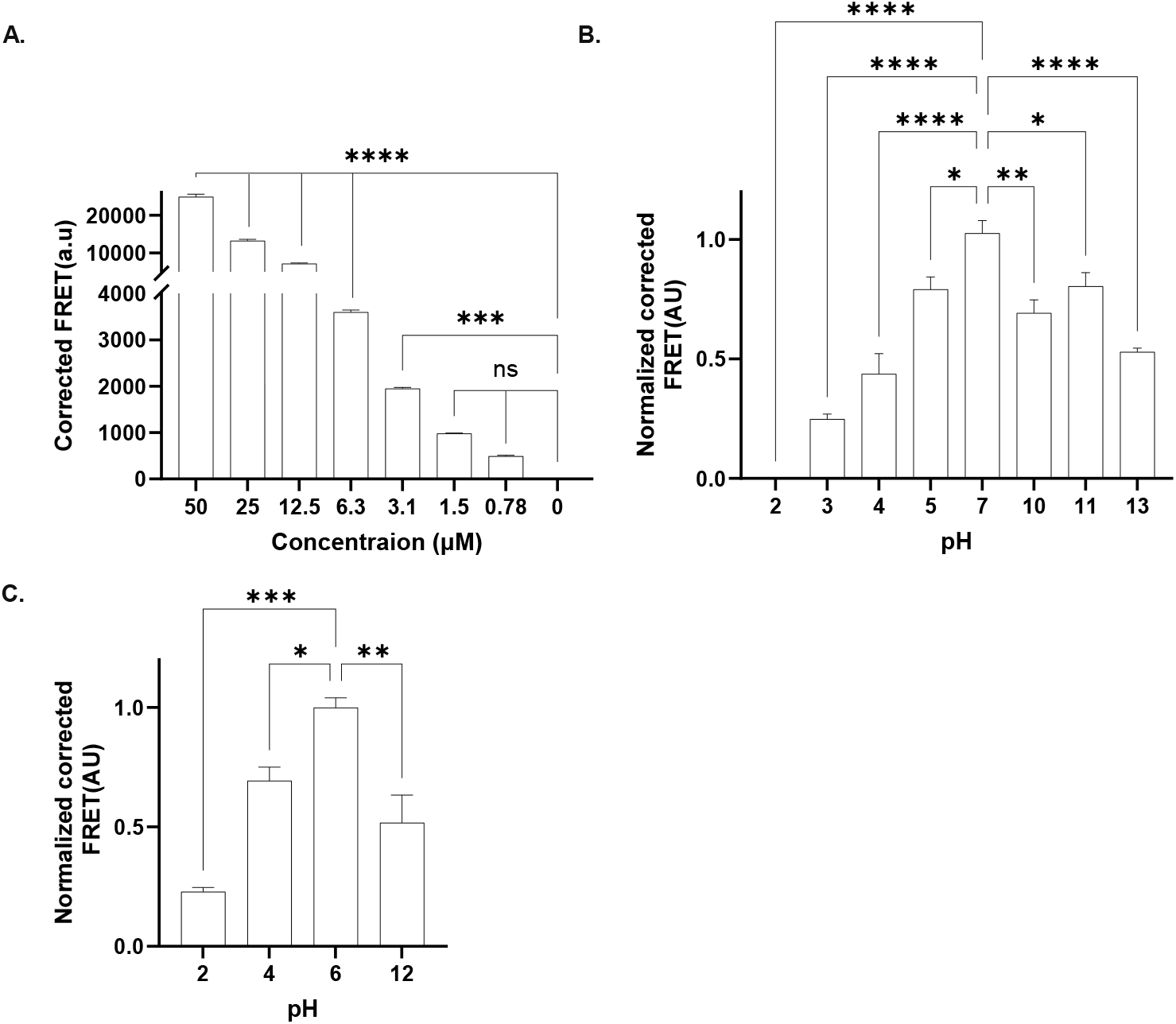
FRET response of the mVenus–Tolles biosensor to pH changes in buffer and food. **(A)** FRET signal of the biosensor measured at increasing concentrations in PBS. **(B)** FRET signal of the biosensor in buffered solutions with variable pH. **(C)** FRET signal response of the biosensor in hummus samples adjusted to defined pH values, demonstrating a comparable pattern to buffer conditions. All FRET readouts show the normalized corrected FRET. Error bars represent the mean ±SD signal of three replicate wells. Statistical significance: ns -non significant; *p < 0.05; **p < 0.001; ***p < 0.005; ****p < 0.0001.

### 3.4 Monitoring food spoilage via mVenus-Tolles FRET response

Following the controlled pH experiments, we next tested the biosensor’s ability to detect spoilage in real food undergoing natural microbial degradation. Hummus paste was placed into a 6-well plate and left exposed to ambient conditions (25 °C). The mVenus–Tolles protein was added to aliquots of the aging hummus, and the FRET signal was recorded. To validate that the FRET changes reflect spoilage due to microbial growth, we simultaneously measured the pH using litmus paper (Figure S2) and quantified bacterial load (CFU/mL) on agar plates each day. As shown in Figures 4A–C, the FRET signal gradually declined coinciding with a drop in pH and a bacterial load.

**Figure 4.**
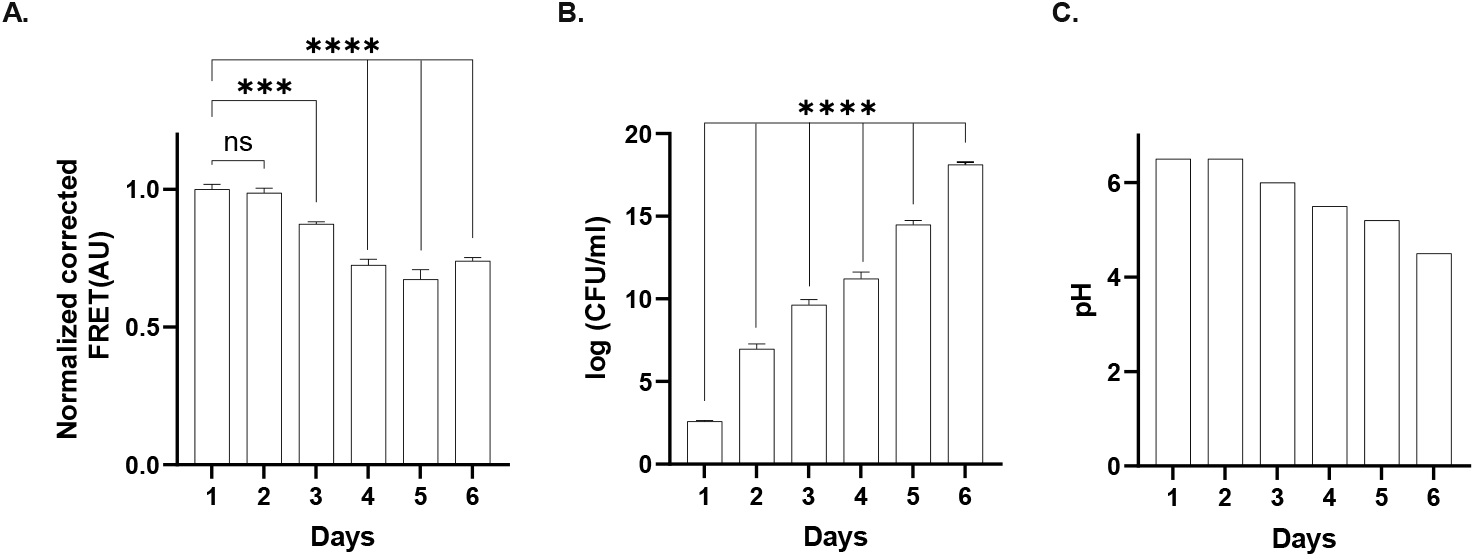
Monitoring spoilage progression in hummus using the mVenus–Tolles FRET biosensor. **(A)** Daily FRET signal originating from fresh hummus samples exposed to ambient conditions (25 °C). **(B)** Corresponding daily pH measurements evaluated using litmus paper. **(C)** Bacterial load as log(CFU/mL) measured via serial dilution colony counting on agar plate. All FRET readouts show the normalized corrected FRET. Errors bars represent the mean ±SD signal of three replicate wells. Statistical significance: ns - non significant; *p < 0.05;**p < 0.001; ***p < 0.005; ****p < 0.0001.

### 3.5 Real-time monitoring of food spoilage via mVenus-Tolles FRET response

To study if the biosensor could be used beyond daily snapshots, we assessed the ability of the mVenus-Tolles biosensor to capture real-time kinetics of pH change. To this end, we conducted continuous FRET measurements in a microplate format. Freshly prepared hummus was aliquoted into a 96-well plate (200 μL per well) and inoculated with 3 μL of a bacterial culture derived from spoiled hummus, pre-grown to an optical density of OD=0.6. Each well was mixed with 5 μM of the mVenus–Tolles biosensor protein. The plate was incubated at 37 °C in a plate reader, and FRET signals were recorded at 60-minute intervals over a 48-hour period. As a control, additional wells were supplemented with a cocktail of ampicillin and chloramphenicol at final concentrations of 100 μg/ml and 32 μg/ml, respectively, to inhibit microbial growth. As shown in **Figure 5**, a sharp decline in the FRET signal was observed in untreated samples, while the antibiotic-treated wells maintained a relatively stable signal. These results confirm that the biosensor can track spoilage kinetics with high temporal resolution, enabling early and automated detection of freshness loss before visible cues emerge.

**Figure 5.**
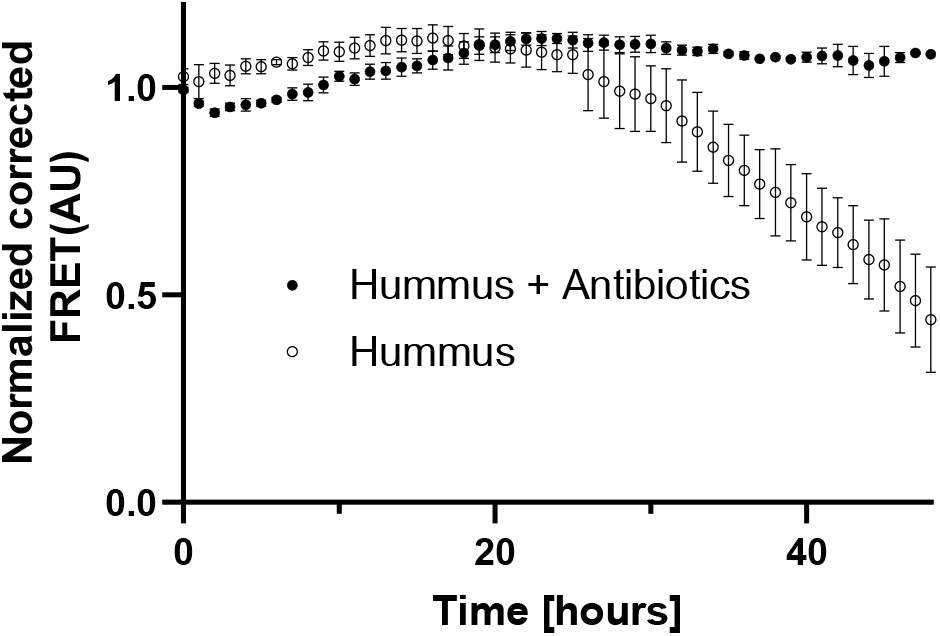
Real-time monitoring of hummus spoilage using the mVenus–Tolles FRET biosensor. Normalized FRET signal over 48 hours in hummus samples incubated at 37 °C. Samples without antibiotics (white) show a sharp decline in FRET signal due to active microbial growth, while samples treated with antibiotics (black) maintain a stable signal. Error bars represent the mean ± SD of three replicate wells.

## 4. Discussion

In this study, we demonstrate the feasibility of using the mVenus–Tolles pHlameleon as a protein-based FRET biosensor for non-invasive monitoring of pH changes associated with food spoilage. By incorporating an N-terminal VNP solubility tag, we achieved high-yield soluble expression of the chimeric protein in E. coli (Figures 1–2), overcoming the typical issue of inclusion body formation when expressing large chimera proteins. The purified biosensor displayed a pH-responsive FRET signal within the pH range of 4–7 (Figure 3), highly relevant for monitoring microbial activity in food systems. When applied to a representative food matrix (hummus), the sensor successfully tracked spoilage both static (single-timepoint) and dynamic (real-time) conditions. Furthermore, the FRET signal correlated with independent measurements of pH and bacterial growth, showing its potential as biosensor of freshness (Figures 4–5).

To our knowledge, this is the first time a recombinant pHlameleon FRET fusion protein applied within a food matrix to monitor pH. Protein-based sensors offer distinct advantages over synthetic or small-molecule alternatives. Proteins are intrinsically biocompatible and, in principle, could be rendered edible, enabling safe incorporation into food products without the risks associated with synthetic dyes or toxic residues. This aligns with recent efforts to use edible materials in advanced optical technologies, such as biolasers [47]. Additionally, fluorescent proteins are highly modular, allowing for the tuning of spectral properties and pKa values to match specific sensing environments. Their recombinant expression in microbial systems also supports sustainable and scalable production which is an important factor for the food industry. Protein biosensors could be multiplexed with other molecular indicators, e.g., for amines or oxygen, to form an integrated “spoilage barcode,” providing a more comprehensive and accurate profile of microbial activity and food degradation leading to waste.

Despite their advantages, protein-based biosensors have several key challenges. Their structural stability can be compromised by environmental factors such as elevated temperatures, endogenous proteases in food, and interactions with complex matrix components. Proper folding and the inclusion of solubility-enhancing tags are often required to prevent aggregation. Moreover, harsh conditions such as low pH or high salt concentrations can alter the spectral properties of fluorescent proteins, potentially affecting sensor performance. The lifetime of such proteins may also be limited, particularly in foods with extended shelf lives. Beyond technical and scientific concerns, the regulatory landscape for using recombinant proteins in food is complex, potentially inhibiting widespread adoption of it. Finally, although lab-scale protein production is relatively inexpensive, the cost-effective purification, formulation, and incorporation of these sensors into consumer-ready food products remains a significant barrier. Overcoming these challenges will require continued innovation and research in the field.

## 5. Conclusion

The study presents the successful application of a recombinant protein-based FRET biosensor, mVenus–Tolles, for real-time, non-invasive pH monitoring in food. The sensor demonstrated robust and sensitive pH-dependent responses within the critical spoilage-related pH range (4–7), accurately correlating with microbial growth in chickpea paste. The use of protein-based biosensors could pave the way for safe and biocompatible alternatives suitable for integration into food and packaging, reducing waste, and enabling intelligent food freshness monitoring technologies.

## CRediT authorship contribution statement

**Matan Gabay:** Methodology, Investigation, Data curation, Writing – original draft, Writing – review & editing. **Marina Sova:** Methodology, Investigation, Project administration. **Tal Laviv:** Conceptualization, Supervision, Writing – review & editing. **Maayan Gal:** Conceptualization, Supervision, Funding acquisition, Writing – original draft, Writing – review & editing.

## Data availability

Data will be made available on request.

