## supplementary information for "Real-Time Sensing of Food Spoilage Using a Recombinant mVenus–Tolles FRET pH Biosensor"

**Plasmid sequences**

**Plasmid#1 – VNP--mVenues-Tolles-Hisx6**

ggggaattgtgagcggataacaattcccctgtagaaataattttgtttaactttaataaggagatataccatgggcagcagccatcaccatcatcaccacagccaggatccgaattcgagctcggcgcgcctgcaggtcgacaagcttgcggccgcataatgcttaagtcgaacagaaagtaatcgtattgtacacggccgcataatcgaaattaatacgactcactataggggaattgtgagcggataacaattccccatcttagtatattagttaagtataagaaggagatatacatatggatgtattcaagaaaggattttcaatagccgatgagggagttgtgggagctgttgagagaGAGAACCTGTACTTTCAGAGCatggtgagcaagggcgaggagctgttcaccggggtggtgcccatcctggtcgagctggacggcgacgtaaacggccacaagttcagcgtgtccggcgagggcgagggcgatgccacctacggcaagctgaccctgaagctGatctgcaccaccggcaagctgcccgtgccctggcccaccctcgtgaccaccctGggctacggcctgcagtgcttcgcccgctaccccgaccacatgaagcagcacgacttcttcaagtccgccatgcccgaaggctacgtccaggagcgcaccatcttcttcaaggacgacggcaactacaagacccgcgccgaggtgaagttcgagggcgacaccctggtgaaccgcatcgagctgaagggcatcgacttcaaggaggacggcaacatcctggggcacaagctggagtacaactacaacagccacaacgtctatatcaccgccgacaagcagaagaacggcatcaaggccaacttcaagatccgccacaacatcgaggacggcggcgtgcagctcgccgaccactaccagcagaacacccccatcggcgacggccccgtgctgctgcccgacaaccactacctgagctaccagtccaagctgagcaaagaccccaacgagaagcgcgatcacatggtcctgctggagttcgtgaccgccgccgggatcactctcggcatggacgagctgtacaaggagctcggtggaggcggttcaggcggaggtggctctggcggtggcggatcgctcgagatggtgagcgtgatcaagcccgagatgaagatcaagctgtgcatgaggggcaccgtgaacggccacaacttcgtgattgagggcgagggcaagggcaacccctacgagggcacccagatcctggacctgaacgtgaccgagggcgcccccctgcccttcgcctacgacatcctgtccacctcgttccagtacggcaacagggccttcaccaagtaccccgccgacatccaggactacttcaagcaggccttccccgagggctaccactgggagaggagcatgacctacgaggaccagggcatctgcaccgccaccagcaacatcagcatgaggggcgactgcttcttctacgacatcaggttcgacggcaccaacttcccccccaacggccccgtgatgcagaagaagaccctgaagtgggagcccgacaccgagaagatgtacgtggaggacggcgtgctgaagggcgacgtgaacatgaggctgctgctggagggcggcggccactacaggtgcgacgtcaagaccacctacaaggccaagaaggaggtgagcctgcccgacgcccacaagatcgaccacaggatcgagatcctggagcacgacaaggactacaacaaggtgaagctgtgcgagaacgccgtggccagggcctccaTGCTGCCCAGCCAGGCCAAGtatagcggaggaagcggaggtacccatcatcatcatcatcattaactcgagtctggtaaagaaaccgctgctgcgaaatttgaacgccagcacatggactcgtctactagcgcagcttaattaacctaggctgctgccaccgctgagcaataactagcataaccccttggggcctctaaacgggtcttgaggggttttttgctgaaacctcaggcatttgagaagcacacggtcacactgcttccggtagtcaataaaccggtaaaccagcaatagacataagcggctatttaacgaccctgccctgaaccgacgacaagctgacgaccgggtctccgcaagtggcacttttcggggaaatgtgcgcggaacccctatttgtttatttttctaaatacattcaaatatgtatccgctcatgaattaattcttagaaaaactcatcgagcatcaaatgaaactgcaatttattcatatcaggattatcaataccatatttttgaaaaagccgtttctgtaatgaaggagaaaactcaccgaggcagttccataggatggcaagatcctggtatcggtctgcgattccgactcgtccaacatcaatacaacctattaatttcccctcgtcaaaaataaggttatcaagtgagaaatcaccatgagtgacgactgaatccggtgagaatggcaaaagtttatgcatttctttccagacttgttcaacaggccagccattacgctcgtcatcaaaatcactcgcatcaaccaaaccgttattcattcgtgattgcgcctgagcgagacgaaatacgcggtcgctgttaaaaggacaattacaaacaggaatcgaatgcaaccggcgcaggaacactgccagcgcatcaacaatattttcacctgaatcaggatattcttctaatacctggaatgctgttttcccggggatcgcagtggtgagtaaccatgcatcatcaggagtacggataaaatgcttgatggtcggaagaggcataaattccgtcagccagtttagtctgaccatctcatctgtaacatcattggcaacgctacctttgccatgtttcagaaacaactctggcgcatcgggcttcccatacaatcgatagattgtcgcacctgattgcccgacattatcgcgagcccatttatacccatataaatcagcatccatgttggaatttaatcgcggcctagagcaagacgtttcccgttgaatatggctcatactcttcctttttcaatattattgaagcatttatcagggttattgtctcatgagcggatacatatttgaatgtatttagaaaaataaacaaataggcatgcagcgctcttccgcttcctcgctcactgactcgctacgctcggtcgttcgactgcggcgagcggtgtcagctcactcaaaagcggtaatacggttatccacagaatcaggggataaagccggaaagaacatgtgagcaaaaagcaaagcaccggaagaagccaacgccgcaggcgtttttccataggctccgcccccctgacgagcatcacaaaaatcgacgctcaagccagaggtggcgaaacccgacaggactataaagataccaggcgtttccccctggaagctccctcgtgcgctctcctgttccgaccctgccgcttaccggatacctgtccgcctttctcccttcgggaagcgtggcgctttctcatagctcacgctgttggtatctcagttcggtgtaggtcgttcgctccaagctgggctgtgtgcacgaaccccccgttcagcccgaccgctgcgccttatccggtaactatcgtcttgagtccaacccggtaagacacgacttatcgccactggcagcagccattggtaactgatttagaggactttgtcttgaagttatgcacctgttaaggctaaactgaaagaacagattttggtgagtgcggtcctccaacccacttaccttggttcaaagagttggtagctcagcgaaccttgagaaaaccaccgttggtagcggtggtttttctttatttatgagatgatgaatcaatcggtctatcaagtcaacgaacagctattccgttactctagatttcagtgcaatttatctcttcaaatgtagcacctgaagtcagccccatacgatataagttgtaattctcatgttagtcatgccccgcgcccaccggaaggagctgactgggttgaaggctctcaagggcatcggtcgagatcccggtgcctaatgagtgagctaacttacattaattgcgttgcgctcactgcccgctttccagtcgggaaacctgtcgtgccagctgcattaatgaatcggccaacgcgcggggagaggcggtttgcgtattgggcgccagggtggtttttcttttcaccagtgagacgggcaacagctgattgcccttcaccgcctggccctgagagagttgcagcaagcggtccacgctggtttgccccagcaggcgaaaatcctgtttgatggtggttaacggcgggatataacatgagctgtcttcggtatcgtcgtatcccactaccgagatgtccgcaccaacgcgcagcccggactcggtaatggcgcgcattgcgcccagcgccatctgatcgttggcaaccagcatcgcagtgggaacgatgccctcattcagcatttgcatggtttgttgaaaaccggacatggcactccagtcgccttcccgttccgctatcggctgaatttgattgcgagtgagatatttatgccagccagccagacgcagacgcgccgagacagaacttaatgggcccgctaacagcgcgatttgctggtgacccaatgcgaccagatgctccacgcccagtcgcgtaccgtcttcatgggagaaaataatactgttgatgggtgtctggtcagagacatcaagaaataacgccggaacattagtgcaggcagcttccacagcaatggcatcctggtcatccagcggatagttaatgatcagcccactgacgcgttgcgcgagaagattgtgcaccgccgctttacaggcttcgacgccgcttcgttctaccatcgacaccaccacgctggcacccagttgatcggcgcgagatttaatcgccgcgacaatttgcgacggcgcgtgcagggccagactggaggtggcaacgccaatcagcaacgactgtttgcccgccagttgttgtgccacgcggttgggaatgtaattcagctccgccatcgccgcttccactttttcccgcgttttcgcagaaacgtggctggcctggttcaccacgcgggaaacggtctgataagagacaccggcatactctgcgacatcgtataacgttactggtttcacattcaccaccctgaattgactctcttccgggcgctatcatgccataccgcgaaaggttttgcgccattcgatggtgtccgggatctcgacgctctcccttatgcgactcctgcattaggaaattaatacgactcactata

**Plasmid#2 – VNP--mVenues-Hisx6**

ggggaattgtgagcggataacaattcccctgtagaaataattttgtttaactttaataaggagatataccatgggcagcagccatcaccatcatcaccacagccaggatccgaattcgagctcggcgcgcctgcaggtcgacaagcttgcggccgcataatgcttaagtcgaacagaaagtaatcgtattgtacacggccgcataatcgaaattaatacgactcactataggggaattgtgagcggataacaattccccatcttagtatattagttaagtataagaaggagatatacatatggatgtattcaagaaaggattttcaatagccgatgagggagttgtgggagctgttgagagaGAGAACCTGTACTTTCAGAGCatggtgagcaagggcgaggagctgttcaccggggtggtgcccatcctggtcgagctggacggcgacgtaaacggccacaagttcagcgtgtccggcgagggcgagggcgatgccacctacggcaagctgaccctgaagctGatctgcaccaccggcaagctgcccgtgccctggcccaccctcgtgaccaccctGggctacggcctgcagtgcttcgcccgctaccccgaccacatgaagcagcacgacttcttcaagtccgccatgcccgaaggctacgtccaggagcgcaccatcttcttcaaggacgacggcaactacaagacccgcgccgaggtgaagttcgagggcgacaccctggtgaaccgcatcgagctgaagggcatcgacttcaaggaggacggcaacatcctggggcacaagctggagtacaactacaacagccacaacgtctatatcaccgccgacaagcagaagaacggcatcaaggccaacttcaagatccgccacaacatcgaggacggcggcgtgcagctcgccgaccactaccagcagaacacccccatcggcgacggccccgtgctgctgcccgacaaccactacctgagctaccagtccaagctgagcaaagaccccaacgagaagcgcgatcacatggtcctgctggagttcgtgaccgccgccgggatcactctcggcatggacgagctgtacaagtatagcggaggaagcggaggtacccatcatcatcatcatcattaactcgagtctggtaaagaaaccgctgctgcgaaatttgaacgccagcacatggactcgtctactagcgcagcttaattaacctaggctgctgccaccgctgagcaataactagcataaccccttggggcctctaaacgggtcttgaggggttttttgctgaaacctcaggcatttgagaagcacacggtcacactgcttccggtagtcaataaaccggtaaaccagcaatagacataagcggctatttaacgaccctgccctgaaccgacgacaagctgacgaccgggtctccgcaagtggcacttttcggggaaatgtgcgcggaacccctatttgtttatttttctaaatacattcaaatatgtatccgctcatgaattaattcttagaaaaactcatcgagcatcaaatgaaactgcaatttattcatatcaggattatcaataccatatttttgaaaaagccgtttctgtaatgaaggagaaaactcaccgaggcagttccataggatggcaagatcctggtatcggtctgcgattccgactcgtccaacatcaatacaacctattaatttcccctcgtcaaaaataaggttatcaagtgagaaatcaccatgagtgacgactgaatccggtgagaatggcaaaagtttatgcatttctttccagacttgttcaacaggccagccattacgctcgtcatcaaaatcactcgcatcaaccaaaccgttattcattcgtgattgcgcctgagcgagacgaaatacgcggtcgctgttaaaaggacaattacaaacaggaatcgaatgcaaccggcgcaggaacactgccagcgcatcaacaatattttcacctgaatcaggatattcttctaatacctggaatgctgttttcccggggatcgcagtggtgagtaaccatgcatcatcaggagtacggataaaatgcttgatggtcggaagaggcataaattccgtcagccagtttagtctgaccatctcatctgtaacatcattggcaacgctacctttgccatgtttcagaaacaactctggcgcatcgggcttcccatacaatcgatagattgtcgcacctgattgcccgacattatcgcgagcccatttatacccatataaatcagcatccatgttggaatttaatcgcggcctagagcaagacgtttcccgttgaatatggctcatactcttcctttttcaatattattgaagcatttatcagggttattgtctcatgagcggatacatatttgaatgtatttagaaaaataaacaaataggcatgcagcgctcttccgcttcctcgctcactgactcgctacgctcggtcgttcgactgcggcgagcggtgtcagctcactcaaaagcggtaatacggttatccacagaatcaggggataaagccggaaagaacatgtgagcaaaaagcaaagcaccggaagaagccaacgccgcaggcgtttttccataggctccgcccccctgacgagcatcacaaaaatcgacgctcaagccagaggtggcgaaacccgacaggactataaagataccaggcgtttccccctggaagctccctcgtgcgctctcctgttccgaccctgccgcttaccggatacctgtccgcctttctcccttcgggaagcgtggcgctttctcatagctcacgctgttggtatctcagttcggtgtaggtcgttcgctccaagctgggctgtgtgcacgaaccccccgttcagcccgaccgctgcgccttatccggtaactatcgtcttgagtccaacccggtaagacacgacttatcgccactggcagcagccattggtaactgatttagaggactttgtcttgaagttatgcacctgttaaggctaaactgaaagaacagattttggtgagtgcggtcctccaacccacttaccttggttcaaagagttggtagctcagcgaaccttgagaaaaccaccgttggtagcggtggtttttctttatttatgagatgatgaatcaatcggtctatcaagtcaacgaacagctattccgttactctagatttcagtgcaatttatctcttcaaatgtagcacctgaagtcagccccatacgatataagttgtaattctcatgttagtcatgccccgcgcccaccggaaggagctgactgggttgaaggctctcaagggcatcggtcgagatcccggtgcctaatgagtgagctaacttacattaattgcgttgcgctcactgcccgctttccagtcgggaaacctgtcgtgccagctgcattaatgaatcggccaacgcgcggggagaggcggtttgcgtattgggcgccagggtggtttttcttttcaccagtgagacgggcaacagctgattgcccttcaccgcctggccctgagagagttgcagcaagcggtccacgctggtttgccccagcaggcgaaaatcctgtttgatggtggttaacggcgggatataacatgagctgtcttcggtatcgtcgtatcccactaccgagatgtccgcaccaacgcgcagcccggactcggtaatggcgcgcattgcgcccagcgccatctgatcgttggcaaccagcatcgcagtgggaacgatgccctcattcagcatttgcatggtttgttgaaaaccggacatggcactccagtcgccttcccgttccgctatcggctgaatttgattgcgagtgagatatttatgccagccagccagacgcagacgcgccgagacagaacttaatgggcccgctaacagcgcgatttgctggtgacccaatgcgaccagatgctccacgcccagtcgcgtaccgtcttcatgggagaaaataatactgttgatgggtgtctggtcagagacatcaagaaataacgccggaacattagtgcaggcagcttccacagcaatggcatcctggtcatccagcggatagttaatgatcagcccactgacgcgttgcgcgagaagattgtgcaccgccgctttacaggcttcgacgccgcttcgttctaccatcgacaccaccacgctggcacccagttgatcggcgcgagatttaatcgccgcgacaatttgcgacggcgcgtgcagggccagactggaggtggcaacgccaatcagcaacgactgtttgcccgccagttgttgtgccacgcggttgggaatgtaattcagctccgccatcgccgcttccactttttcccgcgttttcgcagaaacgtggctggcctggttcaccacgcgggaaacggtctgataagagacaccggcatactctgcgacatcgtataacgttactggtttcacattcaccaccctgaattgactctcttccgggcgctatcatgccataccgcgaaaggttttgcgccattcgatggtgtccgggatctcgacgctctcccttatgcgactcctgcattaggaaattaatacgactcactata

**Plasmid#3 – VNP-Tolles-Hisx6**

ggggaattgtgagcggataacaattcccctgtagaaataattttgtttaactttaataaggagatataccatgggcagcagccatcaccatcatcaccacagccaggatccgaattcgagctcggcgcgcctgcaggtcgacaagcttgcggccgcataatgcttaagtcgaacagaaagtaatcgtattgtacacggccgcataatcgaaattaatacgactcactataggggaattgtgagcggataacaattccccatcttagtatattagttaagtataagaaggagatatacatatggatgtattcaagaaaggattttcaatagccgatgagggagttgtgggagctgttgagagaATGGTGAGCGTGATCAAGCCCGagatgaagatcaagctgtgcatgaggggcaccgtgaacggccacaacttcgtgattgagggcgagggcaagggcaacccctacgagggcacccagatcctggacctgaacgtgaccgagggcgcccccctgcccttcgcctacgacatcctgtccacctcgttccagtacggcaacagggccttcaccaagtaccccgccgacatccaggactacttcaagcaggccttccccgagggctaccactgggagaggagcatgacctacgaggaccagggcatctgcaccgccaccagcaacatcagcatgaggggcgactgcttcttctacgacatcaggttcgacggcaccaacttcccccccaacggccccgtgatgcagaagaagaccctgaagtgggagcccgacaccgagaagatgtacgtggaggacggcgtgctgaagggcgacgtgaacatgaggctgctgctggagggcggcggccactacaggtgcgacgtcaagaccacctacaaggccaagaaggaggtgagcctgcccgacgcccacaagatcgaccacaggatcgagatcctggagcacgacaaggactacaacaaggtgaagctgtgcgagaacgccgtggccagggcctccaTGCTGCCCAGCCAGGCCAAGtatagcggaggaagcggaggtacccatcatcatcatcatcattaactcgagtctggtaaagaaaccgctgctgcgaaatttgaacgccagcacatggactcgtctactagcgcagcttaattaacctaggctgctgccaccgctgagcaataactagcataaccccttggggcctctaaacgggtcttgaggggttttttgctgaaacctcaggcatttgagaagcacacggtcacactgcttccggtagtcaataaaccggtaaaccagcaatagacataagcggctatttaacgaccctgccctgaaccgacgacaagctgacgaccgggtctccgcaagtggcacttttcggggaaatgtgcgcggaacccctatttgtttatttttctaaatacattcaaatatgtatccgctcatgaattaattcttagaaaaactcatcgagcatcaaatgaaactgcaatttattcatatcaggattatcaataccatatttttgaaaaagccgtttctgtaatgaaggagaaaactcaccgaggcagttccataggatggcaagatcctggtatcggtctgcgattccgactcgtccaacatcaatacaacctattaatttcccctcgtcaaaaataaggttatcaagtgagaaatcaccatgagtgacgactgaatccggtgagaatggcaaaagtttatgcatttctttccagacttgttcaacaggccagccattacgctcgtcatcaaaatcactcgcatcaaccaaaccgttattcattcgtgattgcgcctgagcgagacgaaatacgcggtcgctgttaaaaggacaattacaaacaggaatcgaatgcaaccggcgcaggaacactgccagcgcatcaacaatattttcacctgaatcaggatattcttctaatacctggaatgctgttttcccggggatcgcagtggtgagtaaccatgcatcatcaggagtacggataaaatgcttgatggtcggaagaggcataaattccgtcagccagtttagtctgaccatctcatctgtaacatcattggcaacgctacctttgccatgtttcagaaacaactctggcgcatcgggcttcccatacaatcgatagattgtcgcacctgattgcccgacattatcgcgagcccatttatacccatataaatcagcatccatgttggaatttaatcgcggcctagagcaagacgtttcccgttgaatatggctcatactcttcctttttcaatattattgaagcatttatcagggttattgtctcatgagcggatacatatttgaatgtatttagaaaaataaacaaataggcatgcagcgctcttccgcttcctcgctcactgactcgctacgctcggtcgttcgactgcggcgagcggtgtcagctcactcaaaagcggtaatacggttatccacagaatcaggggataaagccggaaagaacatgtgagcaaaaagcaaagcaccggaagaagccaacgccgcaggcgtttttccataggctccgcccccctgacgagcatcacaaaaatcgacgctcaagccagaggtggcgaaacccgacaggactataaagataccaggcgtttccccctggaagctccctcgtgcgctctcctgttccgaccctgccgcttaccggatacctgtccgcctttctcccttcgggaagcgtggcgctttctcatagctcacgctgttggtatctcagttcggtgtaggtcgttcgctccaagctgggctgtgtgcacgaaccccccgttcagcccgaccgctgcgccttatccggtaactatcgtcttgagtccaacccggtaagacacgacttatcgccactggcagcagccattggtaactgatttagaggactttgtcttgaagttatgcacctgttaaggctaaactgaaagaacagattttggtgagtgcggtcctccaacccacttaccttggttcaaagagttggtagctcagcgaaccttgagaaaaccaccgttggtagcggtggtttttctttatttatgagatgatgaatcaatcggtctatcaagtcaacgaacagctattccgttactctagatttcagtgcaatttatctcttcaaatgtagcacctgaagtcagccccatacgatataagttgtaattctcatgttagtcatgccccgcgcccaccggaaggagctgactgggttgaaggctctcaagggcatcggtcgagatcccggtgcctaatgagtgagctaacttacattaattgcgttgcgctcactgcccgctttccagtcgggaaacctgtcgtgccagctgcattaatgaatcggccaacgcgcggggagaggcggtttgcgtattgggcgccagggtggtttttcttttcaccagtgagacgggcaacagctgattgcccttcaccgcctggccctgagagagttgcagcaagcggtccacgctggtttgccccagcaggcgaaaatcctgtttgatggtggttaacggcgggatataacatgagctgtcttcggtatcgtcgtatcccactaccgagatgtccgcaccaacgcgcagcccggactcggtaatggcgcgcattgcgcccagcgccatctgatcgttggcaaccagcatcgcagtgggaacgatgccctcattcagcatttgcatggtttgttgaaaaccggacatggcactccagtcgccttcccgttccgctatcggctgaatttgattgcgagtgagatatttatgccagccagccagacgcagacgcgccgagacagaacttaatgggcccgctaacagcgcgatttgctggtgacccaatgcgaccagatgctccacgcccagtcgcgtaccgtcttcatgggagaaaataatactgttgatgggtgtctggtcagagacatcaagaaataacgccggaacattagtgcaggcagcttccacagcaatggcatcctggtcatccagcggatagttaatgatcagcccactgacgcgttgcgcgagaagattgtgcaccgccgctttacaggcttcgacgccgcttcgttctaccatcgacaccaccacgctggcacccagttgatcggcgcgagatttaatcgccgcgacaatttgcgacggcgcgtgcagggccagactggaggtggcaacgccaatcagcaacgactgtttgcccgccagttgttgtgccacgcggttgggaatgtaattcagctccgccatcgccgcttccactttttcccgcgttttcgcagaaacgtggctggcctggttcaccacgcgggaaacggtctgataagagacaccggcatactctgcgacatcgtataacgttactggtttcacattcaccaccctgaattgactctcttccgggcgctatcatgccataccgcgaaaggttttgcgccattcgatggtgtccgggatctcgacgctctcccttatgcgactcctgcattaggaaattaatacgactcactata

| **Primer Name** | **Sequence** |
| --- | --- |
| PM279_MGp64_BB_Fw  (19-mer) | gagctcggtggaggcggtt |
| PM261_pET22b_MBP_BB_Rev  (26-mer): | GAATTCAGTCTGCGCGTCTTTCAGGG |
| PM280_MGp64_Ins_Gib_Fw  (46-mer) | CCCTGAAAGACGCGCAGACTGAATTCatggtgagcaagggcgagga |
| PM281_MGp64_Ins_Gib_Rev  (42-mer) | aaccgcctccaccgagctccttgtacagctcgtccatgccga |
| VNP15_Gib_open_F  (21-mer) | tagcggaggaagcggaggtac |
| VNP15_open_R  (23-mer): | tctctcaacagctcccacaactc |
| mVenus_Gib_VNP15_F  (49-mer) | gagttgtgggagctgttgagagaatggtgagcaagggcgaggagctgtt |
| TOLLES_Gib_VNP15_R  (43-mer) | gtacctccgcttcctccgctataCTTGGCCTGGCTGGGCAGCA |
| PM290_VNP15_Rev_tag  (23-mer) | tatagcggaggaagcggaggtac |
| P006_P2_rv  (24-mer) | cttgtacagctcgtccatgccgag |
| VNP15_open_R  (23-mer) | tctctcaacagctcccacaactc |
| FOR_TOLLES  (22-mer) | ATGGTGAGCGTGATCAAGCCCG |
